# The gH/gL/gO Complex of Elephant Endotheliotropic Herpesvirus 1A Functions as a Receptor-Binding Complex

**DOI:** 10.64898/2025.12.03.692000

**Authors:** Oliver Debski-Antoniak, Lukas Ermers, Jasper Kroes, Ieva Drulyte, Siu Yi Lee, Victor P.M.G. Rutten, Willem Schaftenaar, Frank J.M. van Kuppeveld, Berend-Jan Bosch, Cornelis A.M. de Haan, Daniel L. Hurdiss, Tabitha E. Hoornweg

**Author notes:** Corresponding authors: Daniel L. Hurdiss; Tabitha E. Hoornweg.

## Abstract

Elephant endotheliotropic herpesvirus (EEHV) causes a fatal haemorrhagic disease in young elephants and represents a major threat to the survival of these endangered species. Despite its clinical importance, the mechanisms of EEHV entry and host cell recognition remain poorly understood. Here, we show that the EEHV1A homolog of the *Betaherpesvirus* glycoprotein O (gO) assembles with glycoproteins H and L (gH/gL) to form a heterotrimeric complex analogous to the human cytomegalovirus (HCMV) gH/gL/gO trimer. Using single-particle cryo-electron microscopy, we determined the structure of the EEHV1A gH/gL/gO complex at 3.2 Å resolution, revealing a conserved overall architecture but distinct gO topology and disulfide arrangement. The EEHV1A trimer, but not gH/gL alone, bound primary elephant endothelial cells, monocytes, and lymphocytes, indicating that gH/gL/gO serves as the receptor-binding complex of EEHV1A. This binding was specifically blocked by sera from naturally infected elephants, confirming its antigenic relevance. Finally, negative-stain electron microscopy polyclonal epitope mapping of gH/gL/gO–Fab complexes identified a Fab population that bound gO near the apex of the gH/gL/gO trimer, suggesting that these antibodies may interfere with gH/gL/gO–receptor interactions. Together, these findings define the molecular architecture and receptor-binding properties of the EEHV1A gH/gL/gO complex and provide a structural framework for the development of vaccines and antibody-based interventions against EEHV-associated disease.

## Introduction

Herpesviruses (order *Herpesvirales*) are large, enveloped double-stranded DNA viruses with genomes ranging from approximately 120 to 240 kilobase pairs (kbp), encoding up to 200 genes (1–3). These viruses infect a wide range of animal species and typically establish lifelong, latent infections within a single or closely related host species (1, 2). The family *Orthoherpesviridae*, which includes all mammalian herpesviruses, is subdivided into the subfamilies *Alphaherpesvirinae*, *Betaherpesvirinae*, and *Gammaherpesvirinae*. Representative human pathogens from each group include Herpes simplex virus 1 (HSV1), Human cytomegalovirus (HCMV), and Epstein–Barr virus (EBV), respectively (1).

Elephant endotheliotropic herpesviruses (EEHVs) are a distinct group of elephant-specific herpesviruses, currently comprising seven recognized species (EEHV1–7), some of which include two subspecies (designated A and B). EEHV species 1, 4, and 5 naturally infect Asian elephants (*Elephas maximus*), whereas EEHV2, -3, -6, and -7 are associated with African elephants (*Loxodonta africana* and *Loxodonta cyclotis*) (4). Virtually all adult elephants, both in managed populations and in range countries, are seropositive for multiple EEHV species, suggesting that elephants are commonly infected with these viruses during their lifetime (5–8). Importantly, young elephants with low or undetectable antibody levels to a given (sub)species are susceptible to an acute and often fatal haemorrhagic disease, known as EEHV haemorrhagic disease (EEHV-HD), when infected with that (sub)species (5, 7, 8). EEHV-HD is the leading cause of death in young elephants under human care, responsible for >50% of all juvenile Asian elephant fatalities in Zoos, and it also affects both managed and free-ranging populations in range countries (4, 9–15).

All EEHV species are phylogenetically distinct from other known herpesviruses but are currently classified within the subfamily *Betaherpesvirinae*, as they share the greatest similarity in gene content and genome organization with members of this group, particularly the roseoloviruses (16, 17). EEHV genomes range from 180 to 205 kbp in length and encode approximately 115–120 genes. Of these, 35 are core herpesvirus genes conserved across all herpesvirus families. Eight genes show homology to orthologs found within the *Betaherpesvirinae*, seven show homology to genes shared by both the *Beta-* and *Gammaherpesvirinae*, and three show homology to genes found in the *Alpha-* and *Gammaherpesvirinae* but not previously identified in *Betaherpesvirinae*. Notably, more than 60 EEHV genes appear unique, with no detectable homologs in other herpesviruses (17, 18). This limited homology in both gene content and genome conservation has prompted proposals to classify EEHVs within a new subfamily, the *Deltaherpesvirinae* (18, 19), although their definitive taxonomic placement remains under discussion.

Entry of herpesviruses into host cells is a multistep process that involves several viral glycoproteins and remains only partially understood. The minimal fusion machinery comprises glycoproteins B, H, and L (gB, gH, and gL), which are conserved across all herpesviruses (20, 21). The class III fusion protein gB forms a homotrimer and functions as the membrane fusogen, while gH and gL form a heterodimeric complex (gH/gL) thought to regulate the fusion process through transient interactions with gB. Beyond its role in fusion regulation, the gH/gL complex also mediates receptor binding in many herpesviruses. In some species, such as the alphaherpesvirus Varicella-zoster virus, gH/gL alone is sufficient for receptor engagement, whereas in others (e.g., HSV1, EBV and HCMV), gH/gL functions in conjunction with accessory viral proteins that may form stable, higher-order complexes. Furthermore, several herpesviruses, including HCMV and EBV, assemble multiple, mutually exclusive gH/gL-based entry complexes that mediate attachment to and entry into distinct host cell types (20).

*In vivo*, EEHVs have been detected in a wide range of tissues, where they infect endothelial cells lining the microvasculature (22, 23). Despite extensive efforts using primary elephant cells, including endothelial cell cultures, *in vitro* propagation of EEHVs has not been achieved (24), limiting our understanding of viral entry and membrane fusion mechanisms. Like all herpesviruses, EEHVs encode gB and gH/gL, which constitute the conserved core fusion machinery. A candidate auxiliary factor is the putative gO protein, which is encoded in all EEHV genomes. In several *betaherpesviruses*, including HCMV, the roseoloviruses, and guinea pig cytomegalovirus, gO forms a heterotrimeric complex with gH and gL, known as gH/gL/gO (25–28). In HCMV, this complex is essential for infection and mediates binding to the cellular receptor (29–32). Although EEHV gO shares limited primary sequence identity with canonical *betaherpesvirus* gO orthologs and is substantially shorter (213 amino acids in EEHV1A compared to 313, 472, 651, and 738 amino acids in HHV7, HCMV (HHV5), HHV6A, and HHV6B, respectively), its proposed function is inferred from its genomic position and similar numbers of cysteine residues, N-glycosylation sites, and isoelectric point (pI) values relative to *betaherpesvirus* gO proteins (16).

Here, we show that EEHV1A gO assembles with gH and gL to form a heterotrimeric complex. We report a cryo-EM structure of the EEHV1A gH/gL/gO complex at 3.2 Å resolution and demonstrate that the trimer, but not gH/gL alone, binds primary elephant endothelial cells, monocytes, and lymphocytes. This binding is specifically blocked by antibodies from naturally infected elephants.

Together, these findings identify EEHV1A gH/gL/gO as the likely receptor-binding complex of EEHV1A and provide a structural framework for future mechanistic and therapeutic studies aimed at controlling EEHV infection and disease.

## Results

### AlphaFold Predictions Support Formation of an EEHV1A gH/gL/gO Heterotrimer

To assess whether EEHV gO or other candidate auxiliary subunits could form a stable ternary complex with the canonical EEHV gH/gL core (5), we performed an expanded AlphaFold3 (AF3) multimer screen (33). In addition to gO, we evaluated several EEHV1A open reading frames (ORF-O, ORF-P, ORF-Q) and two viral envelope proteins (E20A and E50). The HCMV-derived gH, gL and gO homologues were used as a positive control (32). For each test condition, AF3 was run on the following combinations: (1) gH/gL, (2) gH/gL/gO, (3) gH/gL/ORF-O, (4) gH/gL/ORF-P, (5) gH/gL/ORF-Q, (6) gH/gL/ORF-O/P/Q, (7) gH/gL/E20A, (8) gH/gL/E50, and (9) gH/gL/gO from HCMV.

Across all complexes, the HCMV gH/gL/gO prediction produced an ipTM score of 0.80, consistent with its known biological interaction and serving as a benchmark for EEHV. Among the EEHV combinations, only gH/gL and gH/gL/gO produced ipTM scores above this threshold (0.86 for both predictions; Fig. 1A). All other candidate subunits, ORF-O, ORF-P, ORF-Q, E20, and E50, scored below 0.8 in every configuration tested, indicating less confidence for their incorporation into a stable ternary complex with gH/gL (Fig. 1A).

**Figure 1.**
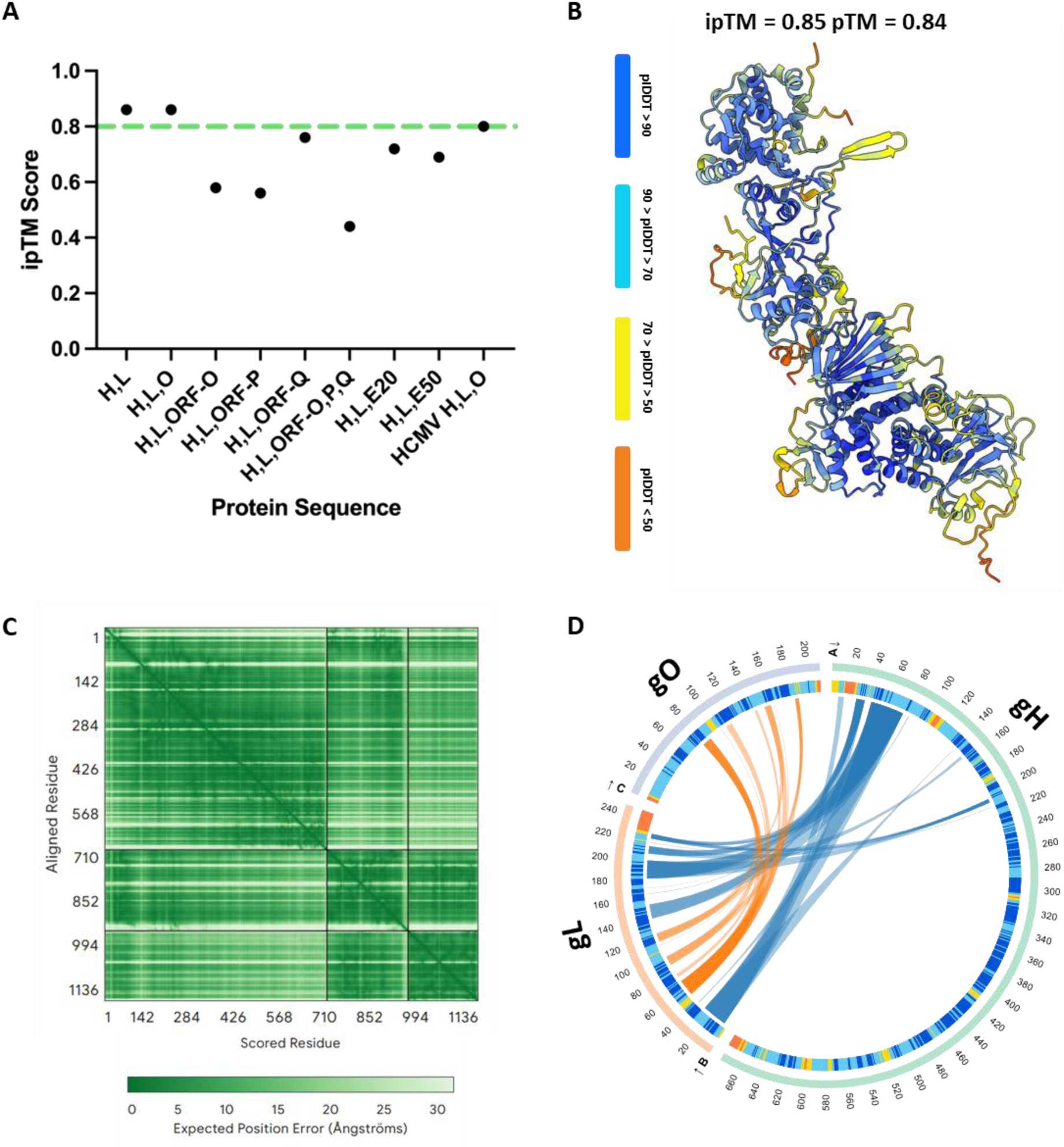
AlphaFold3 prediction and interaction analysis of the EEHV1A gH/gL/gO complex. **(A)** AlphaFold3 (AF3) interaction confidence scores (ipTM) for the EEHV1A gH/gL/gO complex, the gH/gL heterodimer, and all other tested gH/gL–auxiliary protein combinations. **(B)** Top-ranked AF3 model of the EEHV1A gH/gL/gO heterotrimer, coloured by per-residue pLDDT. **(C)** Predicted Aligned Error (PAE) heatmap for the AF3 gH/gL/gO model. **(D)** AlphaBridge circular interaction plot depicting residue–residue interaction communities derived from AF3 confidence metrics.

The AF3 prediction of EEHV1A gH/gL/gO (Fig. 1B) consistently yielded high-confidence geometry and a reproducible ternary complex. As expected, gH and gL were predicted with uniformly high per-residue confidence, with most positions exceeding pLDDT values of 90. The gO chain displayed more variable pLDDT values, including flexible surface-exposed loops with moderate scores, but its structured core and all interface-proximal regions scored above 70, supporting a well-defined interaction surface. The overall architecture closely mirrors the known HCMV gH/gL/gO assembly.

Predicted alignment error (PAE) plots from AF3 (Fig. 1C) further support the integrity of the EEHV gH/gL/gO model. Residues forming the gH–gL and gL–gO interfaces exhibit low expected positional error (typically <5 Å), indicating strong confidence in the relative placement of all three chains. Regions of elevated PAE correspond to flexible termini or loops not involved binding to gL.

To further interpret the AF3 predictions, we analysed the gH/gL/gO model using AlphaBridge (34), a graph-based community clustering tool that reorganizes AF3 confidence metrics (pLDDT, PAE, and predicted distance error) into residue–residue interaction networks. The resulting circular interaction plot (Fig. 1D) revealed a well-defined interaction community linking gO to gL, and a separate community linking gL to gH. These interaction communities correspond to the interface regions highlighted by AF3 as being high-confidence, and no comparable communities were observed in any of the weakly scoring ORF- or envelope-protein models. Collectively, these structural predictions suggest that gO is the only EEHV1A candidate capable of forming a stable and well-supported ternary assembly with gH/gL.

### A disulfide-linked EEHV1A gH/gL/gO heterotrimer forms upon co-expression

We next sought to produce the predicted EEHV1A gH/gL/gO complex in mammalian cells. Single and co-transfections of gH-3×ST, gL-3×ST, and gO-3×ST expression constructs were performed. Following single transfections, proteins of the expected sizes were detected (Fig. 2A; Table S1), and both gL-3×ST and gO-3×ST were found to be secreted efficiently into the supernatant, appearing highly glycosylated. Consistent with previous observations (5), gH-3×ST was not secreted when expressed alone; however, co-expression with gL-3×ST resulted in the production and secretion of both proteins. When gH-3×ST, gL-3×ST, and gO-3×ST were co-expressed, all three proteins were produced and secreted (Fig. 2A). Notably, secretion of gH and gL increased upon co-expression with gO, suggesting that gO may interact with and stabilize the gH/gL heterodimer.

**Figure 2.**
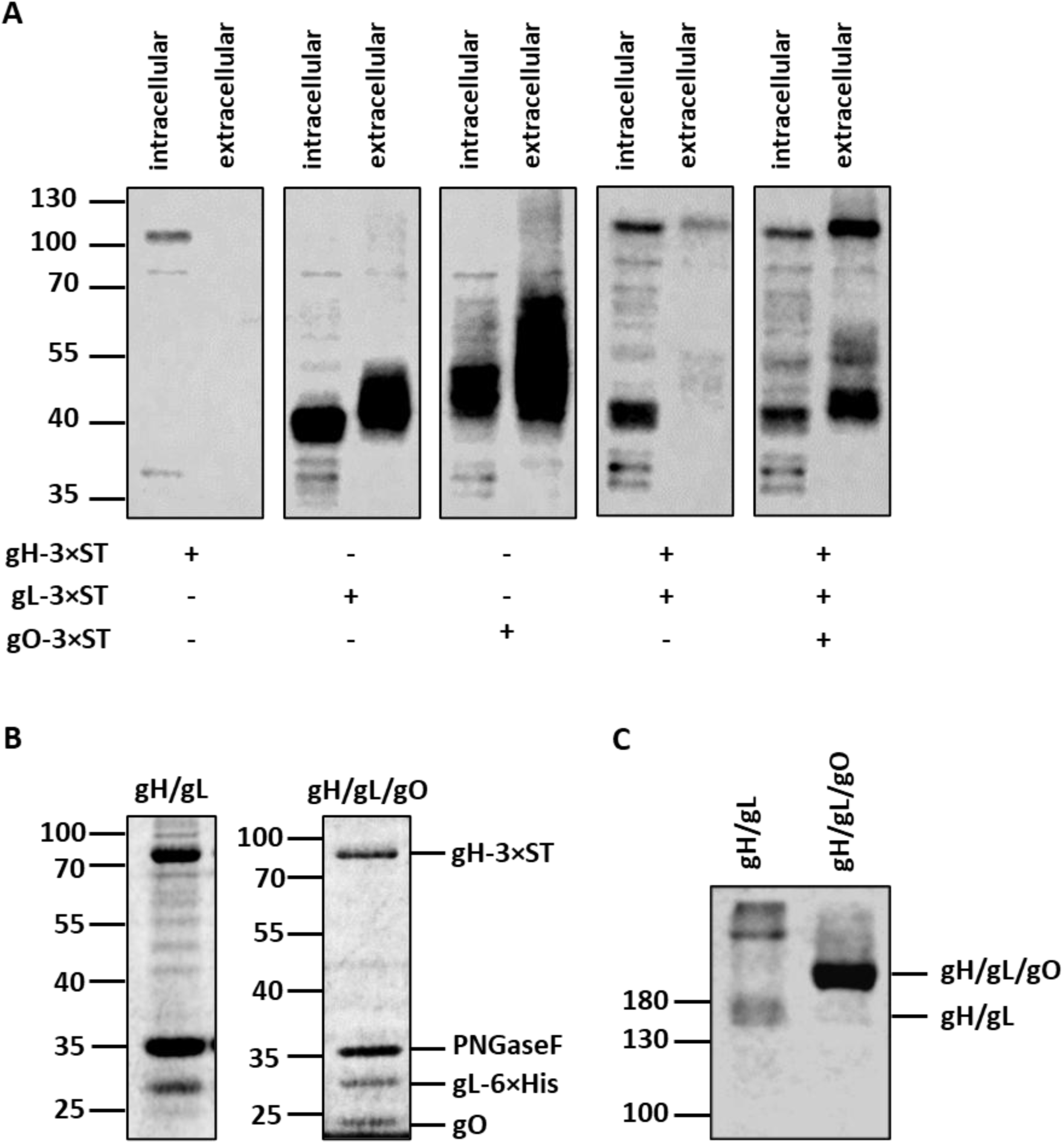
Production and purification of EEHV1A gH/gL/gO. **(A)** Western Blots showing EEHV1A gH-3×ST, gL-3×ST and gO-3×ST production (intracellular) and secretion (extracellular) after single and co-transfections of HEK293T cells. **(B)** Coomassie-stained gels loaded with StrepTactin purified proteins from EEHV1A gH-3×ST + gL-6×His or gH-3×ST + gL-6×His + gO-transfected 293F cells. Samples were deglycosylated by PNGaseF (∼ 35 kDa) and reduced prior to electrophoresis to facilitate analysis. **(C)** Coomassie-stained gels of purified EEHV1A gH-3×ST/gL-6×His or gH-3×ST/gL-6×His/gO separated under non-reducing conditions. In all panels molecular weight markers are indicated on the left side of the gels.

To further investigate whether gH, gL, and gO form a stable protein complex, gH-3×ST and gL-6×His were expressed in the presence or absence of wild-type gO, and the secreted proteins were purified via the StrepTag fused to gH. The Coomassie-stained gels of affinity-purified samples, pretreated with PNGase F to remove N-linked glycans and separated under reducing conditions, are shown in Figure 2B. Co-expression of gH and gL resulted in the formation of a stable heterodimer, as evidenced by the co-purification of gL-6×His with gH-3×ST and the detection of clear protein bands at the expected molecular weights of gH and gL. When gO was co-expressed together with gH and gL, an additional band corresponding to the size of gO (∼24.2 kDa; Table S1) was observed, indicating that gO co-purifies with gH/gL and thus forms a trimeric complex.

In HCMV, disulfide bonds between gH, gL, and gO covalently link the three subunits to form the gH/gL/gO heterotrimer. We previously demonstrated that EEHV gH and gL are likewise covalently connected through conserved cysteines shared with HCMV (5). To determine whether EEHV1A gO is similarly linked to the gH/gL complex by disulfide bonds, secreted gH/gL and gH/gL/gO fractions were analyzed by SDS–PAGE under non-reducing conditions. As expected, co-expression of gH and gL yielded a protein band between 130 and 180 kDa, consistent with the size of a glycosylated gH/gL heterodimer (∼145 kDa) (Fig. 2C). Upon co-expression of gH, gL, and gO, a single higher–molecular-weight band (>180 kDa) was observed, corresponding to the expected size of the glycosylated gH/gL/gO heterotrimer (∼185 kDa). These results confirm the formation of the EEHV1A gH/gL/gO complex and indicate that the constituent proteins are covalently linked via disulfide bonds.

### Structural validation of the EEHV1A gH/gL/gO complex by cryo-EM

To test the validity of the AlphaFold prediction and elucidate the architecture of the gH/gL/gO heterotrimer, we used single-particle cryo-EM. Initial three-dimensional classification revealed one dominant class corresponding to a heterotrimeric assembly, which refined to a global resolution of 3.2 Å (Fig. 3A and S1). The final reconstruction was of high overall quality: gH, gL, and gO were well resolved throughout their core regions, enabling unambiguous backbone tracing and placement of most side chains (Fig. S1).

**Figure 3.**
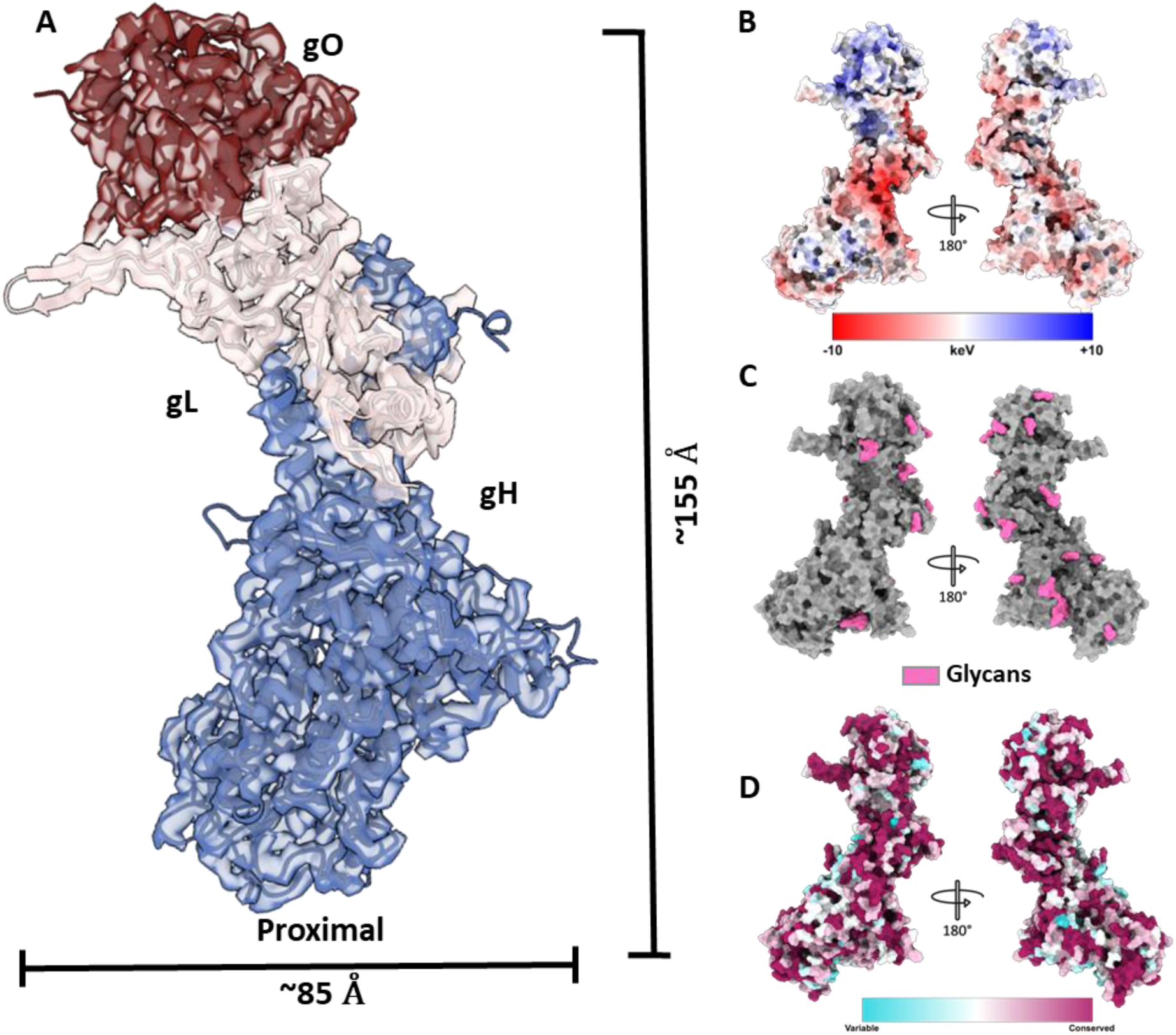
**Structure of EEHV1A heterotrimer gH/gL/gO**. (**A**) Overall cryo-EM map of EEHV1A gH/gL/gO Trimer complex with the fitted atomic model. **(B)** Electrostatic surface of EEHV1A gH/gL/gO Trimer complex **(C)** Glycosylation-site distribution of EEHV1A gH/gL/gO Trimer complex (same views as in **B**). **(D)** Conservation surface of EEHV1A gH/gL/gO Trimer complex (same views as in **B**).

The complex measures approximately 155 × 85 Å and adopts a compact heterotrimeric architecture, in which gH and gL form a contiguous scaffold that engages gO along one face (Fig. 3A and S2). The AlphaFold prediction was used for initial rigid-body fitting, followed by further model adjustment and refinement. Despite the low sequence identity (<20%) between EEHV1A gO and HCMV gO, and the substantially shorter length of the EEHV1A protein, the relative placement of gO adjacent to gL closely mirrors the arrangement observed in HCMV gH/gL/gO trimer structures (Fig. S2; (32)).

The cryo-EM map reveals clearly resolved disulfide linkages that stabilize the gH/gL/gO heterotrimer. In analogy to HCMV, EEHV1A employs gO-C189 to form a disulfide bond with gL-C166 (corresponding to HCMV gO-C351 and gL-C144), and these cysteines are conserved across human betaherpesviruses and all EEHV (sub)species (Fig. S3-S5). Other cysteines stabilizing the gH/gL/gO scaffold, corresponding to HCMV gH-C95 and gL-C47, are likewise conserved in EEHV1A (gH-C78 and gL-C67) and in the roseoloviruses, indicating that the overall disulfide-bonding architecture is broadly conserved (Figs. S5–S6). Next to the conserved cysteines, all EEHV species contain two cysteines not present in gO of the betaherpesviruses, which form an additional disulfide bond within the gO protomer (Fig. S3).

Electrostatic surface analysis revealed a pronounced, surface-exposed electropositive patch on gO that remains accessible on the trimer surface (Fig. 3B and Fig. S7B). Several N-linked glycans were clearly resolved in the cryo-EM map and modelled into the corresponding densities, revealing a glycan distribution that does not occlude this basic, and relatively well conserved, region (Fig. 3C-D and Fig. S7C-D). The accessibility of this electropositive patch is consistent with its potential role as a receptor-binding site within gO.

A focused analysis of gO (Fig. S7) shows that it forms a compact, predominantly α-helical structure that sits within a recessed region of gL, engaging it through a ∼1500 Å² interface (Fig. 3A). The gO–gL contact is stabilized by ten hydrogen bonds and a single intermolecular disulfide linkage. FoldSeek analysis identified HCMV gO as the closest structural homolog, which likewise contains a core cytokine-like fold (Fig. S7E). However, unlike HCMV gO, the EEHV1A protein lacks the additional β-sheet and α-helical extensions that give its HCMV counterpart a more elongated architecture, and consequently adopts a much more compact fold (Fig. S2).

Sequence alignments across betaherpesviruses and EEHV1A reveal that gO proteins are highly variable in both length and composition, yet a small structural core and several conserved cysteines are retained (Fig. S3). The conservation of gO-C189 among EEHV species (Fig. S4), together with its disulfide linkage to gL-C166, suggests that EEHV lineages share a conserved gH/gL/gO heterotrimeric architecture despite extensive sequence divergence in gO (up to ∼70% at the amino acid level). Taken together, the preserved three-chain arrangement and the lineage-specific disulfide connectivity support a conserved functional role for gO as an auxiliary subunit in viral entry across EEHV and other betaherpesviruses.

### Antibodies of naturally EEHV1A-infected elephants are highly reactive to gH/gL/gO

To assess whether the EEHV1A gH/gL/gO complex and its individual subunits are recognized by antibodies from naturally infected elephants, gH/gL, gH/gL/gO, gL, and gO were coated at equimolar concentrations, and serum reactivity against these proteins was measured by ELISA. gH alone could not be included, as this protein is not secreted in the absence of gL (Fig. 2A), precluding its purification. All sera showed strong reactivity to gH/gL and gH/gL/gO, whereas much weaker responses were observed to gL and gO when tested outside the context of the heterotrimer (Fig. 4A). Only a small number of sera exhibited detectable reactivity to isolated gO, but none reacted to gL when presented alone. Increased reactivity to gH/gL/gO compared with gH/gL was consistently observed in sera with non-saturated gH/gL responses (OD < 4, n = 13). For these sera, the gO-specific ΔOD values were compared with the difference in reactivity between gH/gL/gO and gH/gL (ΔOD^gH/gL/gO-gH/gL^; Table S2). Despite the enhanced recognition of gH/gL/gO relative to gH/gL, reactivity to gO alone remained low in most sera. This finding suggests that the majority of gO-targeting antibodies recognize conformational epitopes spanning multiple protomers of the gH/gL/gO complex, or that gO adopts a distinct conformation when expressed outside the context of the heterotrimer.

**Figure 4.**
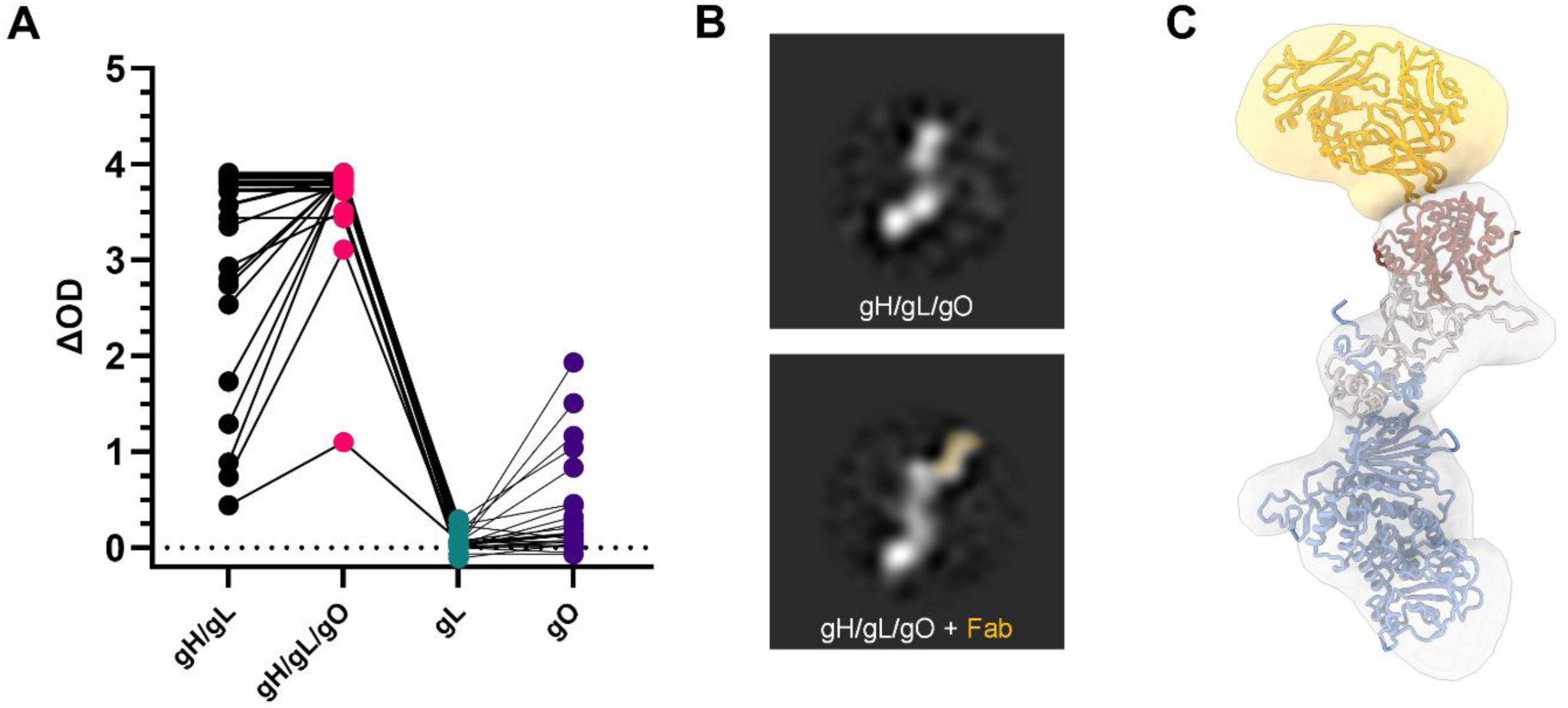
Recognition of EEHV1A gH/gL/gO by antibodies of naturally EEHV1A-infected Asian elephants. **(A)** ΔOD values obtained for 29 sera from 24 individual Asian elephants using gH/gL, gH/gL/gO, gL and gO ELISAs. Per serum sample, the ΔOD values obtained for the different antigens are connected by a line. Concentrations equimolar to 100 ng/well gH/gL were coated for all proteins and sera were tested at a 1:100 dilution. OD = optical density **(B)** Reference-free 2D class averages of the EEHV1A gH/gL/gO complex obtained by negative-stain electron microscopy. The top panel shows the unbound gH/gL/gO trimer, and the bottom panel shows the complex after incubation with the polyclonal Fab mixture. Additional density corresponding to Fab binding is false-coloured for clarity. **(C)** Three-dimensional reconstruction of the gH/gL/gO–Fab complex from negative-stain electron microscopy. The Fab density (orange) is positioned near the apex of the gO subunit, consistent with antibody recognition of an exposed surface on gO. The underlying gH/gL/gO trimer and the rigid-body–fitted Fab fragment (PDB ID: 7FAB) are shown as segmented densities with corresponding fitted models.

To further examine the antibody specificities associated with the enhanced gH/gL/gO response observed, we performed EMPEM using Fab fragments generated from serum of an animal with increased gH/gL/gO- over gH/gL-reactivity. Purified gH/gL/gO was incubated with these Fab fragments, and the resulting mixtures were enriched for immune complexes by size-exclusion chromatography prior to analysis by negative-stain transmission electron microscopy (ns-TEM). Reference-free 2D classification revealed class averages consistent with the gH/gL/gO trimer and putative polyclonal Fab-bound complexes.

Downstream 3D classification yielded a single, stable 3D class that was most consistent with a gH/gL/gO trimer engaged by a Fab positioned along the exposed surface of gO (Fig. 4B). Although the resolution of the negative-stain reconstruction limits precise interpretation, the recovered density suggested Fab interaction with an accessible region of gO near the apex of the trimer (Fig. 4C). While other plausible gH/gL/gO–Fab complexes were apparent in the 2D class averages, these did not produce stable 3D classes, most likely due to the limited particle numbers obtained.

Together, these observations suggest that naturally infected animals can produce an enhanced gH/gL/gO response in which gO-reactive antibodies represent a prominent component, although additional epitope specificities cannot be excluded.

### EEHV1A gH/gL/gO binds elephant but not human cells, and binding is inhibited by heterotrimer-specific antibodies

Because HCMV gH/gL/gO mediates receptor binding to promote viral entry, we investigated whether the EEHV1A gH/gL/gO complex serves a similar function. To this end, EEHV1A gH/gL or gH/gL/gO complexes were coupled to StrepTactin–APC and incubated with elephant umbilical cord endothelial cells (EUVECs), elephant peripheral blood mononuclear cells (PBMCs), and human embryonic kidney (HEK) cells. Cell-associated APC fluorescence was then analyzed by flow cytometry. Elephant PBMCs were further subdivided into monocyte and lymphocyte populations based on forward- and side-scatter profiles. Representative histograms of cell-associated fluorescence after incubation with gH/gL/gO–APC, gH/gL–APC, or uncoupled StrepTactin–APC are shown in Figure 5A. High levels of gH/gL/gO binding were observed for all three elephant cell types, whereas no appreciable binding was detected for human cells (Fig. 5B). Importantly, gH/gL alone showed no significant binding to elephant cells, confirming that cell attachment is specific to the gH/gL/gO heterotrimer.

**Figure 5.**
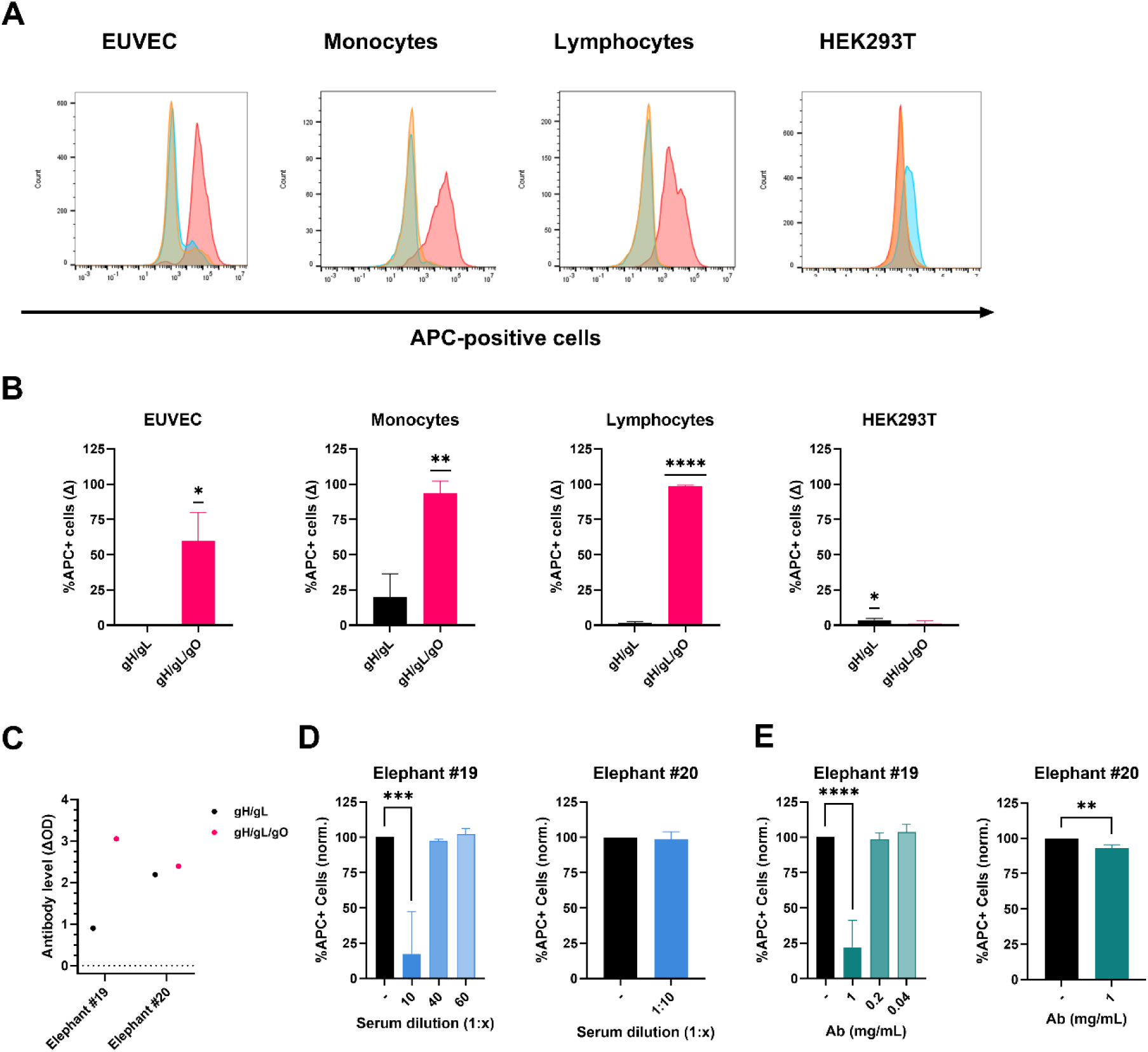
Binding of EEHV1A gH/gL(/gO) to elephant and human cells. **(A)** Representative histograms of gH/gL(/gO) binding to EUVEC, Asian elephant monocytes, Asian elephant lymphocytes and HEK293T. Histograms show the number of cells bound by uncoupled Streptactin-APC (yellow), gH/gL-APC (blue), or gH/gL/gO-APC (red). **(B)** Percentage gH/gL(/gO)-bound EUVEC, monocytes, lymphocytes or HEK293T cells. In all conditions, the percentage APC-positive cells in the uncoupled Streptactin-APC condition was subtracted from the percentage of APC-positive cells in the gH/gl(/gO)-APC condition. Statistical significance was tested by one sample t test, n = 3. **(C)** gH/gL- or gH/gL/gO-specific antibody levels of elephants #19 and #20 as detected by ELISA. Protein concentrations equimolar to 40 ng/well gH/gL were coated and sera were tested at an 1:800 dilution. **(D)** Percentage of gH/gL/gO-bound EUVEC cells after pre-incubation with fourfold dilutions of elephant sera. Values shown were normalized to the percentage of positive cells in the positive control (EUVEC + gH/gL/gO-APC). Statistical significance was tested by one way ANOVA (left panel) or unpaired t-test (right panel), n = 3. **(E)** Percentage of gH/gL/gO bound EUVEC cells after pre-incubation with decreasing concentrations of antibodies isolated from elephant sera. Values shown were normalized and statistical significance was tested as in panel (**D)**, n = 3. Statistical analyses were performed using Graphpad Prism: * = *p* ≤ 0.05, * = *p* ≤ 0.01, *** = *p* ≤ 0.001, **** = *p* ≤ 0.0001.

To verify the specificity of gH/gL/gO binding to elephant cells, the APC-coupled gH/gL/gO complex was preincubated with sera from the same adult elephants as used in Fig. 4B-C prior to incubation with elephant endothelial cells, the primary EEHV target cell *in vivo* (23). At a high serum dilution (1:800), both sera showed strong reactivity to gH/gL/gO; however, serum from elephant #19 displayed greater reactivity to gH/gL/gO than to gH/gL, whereas serum from elephant #20 reacted similarly to both proteins (Fig. 5C). This pattern suggests that antibodies from elephant #19 recognize epitopes spanning gO-containing regions of the trimer, while antibodies from elephant #20 primarily target gH/gL-specific epitopes. When these sera were preincubated with gH/gL/gO prior to cell binding, serum from elephant #19 effectively blocked gH/gL/gO attachment to elephant endothelial cells at high concentrations, whereas serum from elephant #20 did not (Fig. 5D). Similar results were obtained when antibodies purified from these sera were tested (Fig. 5E). Together, these findings demonstrate that gH/gL/gO binds a receptor on elephant cells and that this interaction can be inhibited only by antibodies recognizing gO in the context of the gH/gL/gO heterotrimer.

## Discussion

In the current study, we show that EEHV1A gH, gL and gO form a heterotrimer that functions as the viral receptor binding complex. The molecular structure of the protein complex is reminiscent of gH/gL/gO of the betaherpesvirus HCMV, yet the gO domain that primarily mediates interactions between HCMV gH/gL/gO and its cellular receptor (32) is lacking in EEHV1A gO, suggesting that EEHV gH/gL/gO interacts with its (still to be identified) cellular receptor in a different manner. Importantly, we observed that antibodies in sera of EEHV-infected elephants could block gH/gL/gO-receptor interactions, suggesting that EEHV1A gH/gL/gO is an important target for the development of preventive therapies against EEHV-HD.

EEHV1A gO is considerably shorter than HCMV gO and the sequence similarity between both proteins is less than 20%, yet important protein functions remained conserved. For example, HCMV gO assists in gH/gL(/gO) passage of the ER to the Trans Golgi network (25) and in absence of gO, gH/gL incorporation into the virion is severely hampered (35). Similarly, excretion of EEHV1A gH/gL was substantially enhanced upon co-expression with gO. More importantly, in analogy to HCMV gH/gL/gO, EEHV1A gH/gL/gO, but not gH/gL alone, binds primary elephant cells and therefore likely functions as the viral receptor binding complex. Although HCMV gH/gL/gO is strictly required for infection of all target cells (31, 36), an additional gH/gL-based pentameric complex is used for infection of certain cell types, including endothelial cells and monocytes (29, 30, 37). Whether EEHV may also require additional proteins next to gH/gL/gO to infect its target cells remains to be determined.

Structurally, the EEHV1A gH/gL/gO trimer recapitulates the overall “boot”-shaped architecture first described for HCMV. Both complexes are anchored by the gH transmembrane helix, yet in the EEHV1A trimer a pronounced kink between gH and gL re-orients gL and gO upwards relative to the slightly tilted gH domain; by contrast, the HCMV trimer adopts a less kinked, more rear-leaning geometry. The gO protomers of both EEHV1A and HCMV were found to have compact central domains in which a cytokine-like fold is organized around a small β-sheet scaffold, and this central domain appears to be conserved across all human betaherpesviruses and EEHV (Fig. S3-S4). The most striking molecular difference between HCMV and EEHV1A gO is the absence in EEHV1A gO of the extended N-terminal β-sheet domain (β1–β5) that, in HCMV, contributes substantially to contacts with known receptors such as PDGFRα and TGFβR3 (32). Instead, the N-terminal domain of EEHV1A contains two α-helices. As a result, it is unlikely that EEHV1A engages the elephant orthologues of PDGFRα and TGFβR3 by the same footprint or mechanism. Instead, several features of the EEHV1A trimer point to an alternative mode of recognition: EEHV1A gO is shorter, retains a conserved core, presents a largely electropositive surface that is not extensively masked by glycans, and displays a cytokine-like core fold with helical embellishments—properties that could favour proteinaceous receptors recognizing surface charge or a structurally conserved fold rather than an HCMV-like β-sheet interface.

In line with the structural similarities between EEHV1A and HCMV gH/gL/gO, our observation that EEHV1A gH/gL/gO is well-recognized by sera of naturally EEHV1A-infected elephants indicates the protein is folded in a native manner. Most sera showed clearly increased reactivity to gH/gL/gO as compared to gH/gL, yet reactivity to gO out of context of the trimer was low or absent, suggesting that the isolated subunit does not expose immunodominant epitopes or may be misfolded. Notably, using polyclonal epitope mapping, we identified an elephant Fab fragment that interacted with an accessible region of gO near the apex of the gH/gL/gO trimer. Since the serum sample of the same animal (elephant #19) blocked the interaction between gH/gL/gO and its cellular receptor, a critical first step in virus infection, we presume the detected Fab fragment could potentially be neutralizing. To determine whether the identified Fab fragment can indeed neutralize EEHV1A infection and may thus be of therapeutic interest, a viral neutralization assay needs to be developed.

The production and structural characterization of EEHV1A gH/gL/gO as presented in this study will enable future studies into EEHV1A receptor identification, mode of receptor binding, and antibody-mediated neutralization of EEHV infections. Moreover, gH/gL/gO is considered to be a good candidate antigen for an EEHV vaccine since it is (i) immunogenic, (ii) produced in the native conformation, (iii) serves a critical role early in infection and (iv) since antibodies against gH/gL correlate with protection against EEHV-HD (14). This conclusion together with the highly lethal nature of EEHV1A-HD and the fact that no effective treatment or vaccine for EEHV1A is available to date, prompted us to initiate a safety and efficacy study of an adjuvanted EEHV1A gH/gL/gO subunit vaccine in elephants.

## Materials and Methods

### In-silico Protein–Protein Interaction Screening Using AlphaFold Multimer

Amino acid sequences of all EEHV1A proteins (gH, gL, gO, ORF-O, ORF-P, ORF-Q, E20A and E50; Kimba strain; GenBank accession number KC618527.1) and HCMV gH, gL and gO (Merlin strain; Genbank accession number NC_006273.2) were retrieved from GenBank and models were generated using AlphaFold3 (AF3) multimer (33). Signal peptides and transmembrane regions were removed prior to modelling.

Predicted complexes were evaluated based on ipTM/pTM scores, per-residue pLDDT, and predicted aligned error (PAE) matrices. For reporting, we calculated: (i) model ipTM and pTM scores; (ii) mean per-chain and interface pLDDT (reported as median ± IQR); and (iii) mean PAE across interface residue pairs. Interface residues were defined as those with any atom ≤ 5.0 Å from a residue on the partner chain, and interface buried surface area (BSA) was computed using FreeSASA (probe radius = 1.4 Å).

Models were classified as high-confidence if they satisfied the following thresholds: ipTM ≥ 0.7, mean interface PAE ≤ 5 Å, and interface pLDDT ≥ 70–80 at contact residues.

### Protein production, purification, and analysis

Protein expression constructs for EEHV1A gH-3×ST, gL-3×ST and gL-6xHis were all described before (5, 7). Since EEHV1A gO is predicted to be secreted (SecretomeP; https://services.healthtech.dtu.dk/services/SecretomeP-2.0/), but does not contain a signal peptide (SignalP; https://services.healthtech.dtu.dk/services/SignalP-6.0/), a full length codon-optimized cDNA (accession number AGG16085.1; GenScript) was cloned into a pFRT expression plasmid (Thermo Fisher Scientific, Waltham, MA, USA), either in frame with a C-terminal triple StrepTag (3×ST) or directly followed by a stop codon, as described previously (5, 7).

For analysis of protein production and secretion, individual plasmids or a combination of the gH, gL and/or gO expression plasmids were transfected into HEK293T cells using polyethylenimine (PEI; PolySciences, Warrington, PA, USA) as described previously (5, 7). Five days post transfection, cell culture media were harvested and the cells were lysed using RIPA buffer, cell culture media and lysates were centrifuged, supernatants were collected and subjected to SDS-PAGE, as described previously (5, 7). Proteins were transferred to a PVDF membrane using the Transblot Turbo System (BioRad, Hercules, CA, USA) and subsequently stained using a horseradish peroxidase (HRP)-conjugated monoclonal anti-StrepTag antibody (Iba). Signals were detected by use of enhanced chemiluminescence (ECL) Western blotting substrate (Pierce, Thermo Fisher Scientific) and the Odyssey imaging system (LI-COR).

For protein purification, individual plasmids or combinations of the gH-3×ST, gL-6×His and the gO expression plasmids were transfected into FreeStyle 293-F cells (Thermo Fisher Scientific) using PEI. Five days after transfection, cell culture media were harvested, cleared of debris, after which proteins were purified using Strep-Tactin Sepharose beads (Iba), as described previously (5, 7). The purity and integrity of all proteins were checked, and protein concentrations were estimated using quantitative densitometry on GelCode Blue-stained protein gels (Thermo Fisher Scientific) containing bovine serum albumin (BSA) standards. Gels were imaged and analysed using the Odyssey imaging system (LI-COR). When indicated, protein fractions were deglycosylated by PNGaseF (NEB) prior to gel electrophoresis to facilitate protein analysis and quantification.

### Cryo-Electron Microscopy Sample Preparation and Data Collection

Purified gH/gL/gO complex was thawed on ice, and 3 µL of sample was applied to glow-discharged Quantifoil R1.2/1.3 grids (Quantifoil Micro Tools GmbH) using a Quorum GloQube system (20 mA, 30 s). Grids were blotted for 4 s with a blot force of 0 and plunge-frozen in liquid ethane using a Vitrobot Mark IV (Thermo Fisher Scientific).

Cryo-EM data were collected on a Thermo Scientific Krios G4 Cryo-Transmission Electron Microscope operated at 300 kV, equipped with a Selectris X energy filter and a Falcon 4i direct electron detector operating in Electron Event Representation (EER) mode. To alleviate preferred particle orientation, the stage was tilted by 33° during data collection. A total of 4,504 movies were acquired at a nominal magnification of 165,000×, corresponding to a calibrated pixel size of 0.73 Å/pixel, with defocus values ranging from –0.75 µm to –1.5 µm.

A full list of data collection parameters is provided in Table S3.

### Single-Particle Image Processing

Image processing was performed using the CryoSPARC software package (v.4.4; (38)). After patch motion correction and CTF estimation, particles were automatically picked using the blob picker, extracted with 4.2× binning, and subjected to several rounds of 2D classification. Particles from class averages displaying well-defined high-resolution features were selected for *ab initio* reconstruction into three classes.

Particles corresponding to the complete gH/gL/gO complex were further refined through one round of heterogeneous refinement to remove low-quality particles, followed by re-extraction at 2.5× binning. Three additional rounds of heterogeneous refinement were performed to further clean the dataset. The resulting particles underwent reference-based motion correction and final non-uniform refinement, yielding a reconstruction at 3.2 Å resolution.

A detailed data processing workflow is shown in Figure S8.

### Model Building and Refinement

Model building was performed using UCSF Chimera (v1.15.0; (39)) and Coot (v0.9.6; (40)). An initial model of the gH/gL/gO complex was generated using AlphaFold3 (33).

Individual subunits were rigid-body fitted into the cryo-EM density map using the *Fit in Map* tool in UCSF Chimera, and then combined into a single composite model. The resulting model was manually adjusted in Coot using the *Real-Space Refinement* and *Carbohydrate Module* tools (41), as well as *Sphere Refinement* for local optimization.

To further improve model-to-map agreement, Namdinator (42) was employed to perform molecular dynamics–based flexible fitting. Subsequent iterative cycles of manual adjustment in Coot and real-space refinement in Phenix (43) were conducted to minimize rotamer, bond angle, and Ramachandran outliers. Secondary structure and non-crystallographic symmetry (NCS) restraints were applied throughout Phenix refinement.

The final model was validated in Phenix using MolProbity (44), and glycan geometries were assessed with Privateer (45, 46).

### Structure Analysis and Visualization

Structural analysis and figure preparation were performed using UCSF ChimeraX (47). Interface analysis was carried out with PDBePISA (48), structural homology searches were performed using FoldSeek (49), and AF3-derived confidence metrics were further interpreted using AlphaBridge (34). All structural biology software used in this study was installed and managed through the SBGrid consortium (50).

### Enzyme-Linked Immunosorbent Assay

A total of 29 sera from 24 individual Asian elephants (*Elephas maximus*), aged 0.7 to 53 years, from different European zoological collections were subjected to ELISA. All blood samples were taken aseptically from ear or leg veins by zoo veterinary staff (as part of routine management recommended by the elephant Taxon Advisor Group of the European Association of Zoos and Aquaria), and sera were transported at 4 °C to our institute for diagnostic purposes. Sera were stored at −20 °C prior to use.

ELISAs were essentially performed as previously described for EEHV1A gH/gL (5, 7). To ensure coating of equal amounts of the different proteins, antigen dilution ranges were coated onto ELISA plates, blocked, and probed using the HRP-conjugated monoclonal anti-StrepTag detection antibody (Iba) to determine the amounts of proteins needed to obtain equal OD values. Unless indicated otherwise, equimolar protein concentrations to 100 ng gH/gL were coated per well, sera were tested at a 1:100 dilution, and recombinant protein A/G-HRP (0.5 µg/mL; Pierce) was used as the secondary conjugate to detect the binding of elephant antibodies.

### Electron Microscopy Polyclonal Epitope Mapping

Elephant antibodies were purified by mixing 1 ml serum with 0.5 ml prewashed protein A/G Plus agarose beads (Pierce) in a total volume of 5 ml PBS and incubating the mixture overnight at 4°C on a roller bank. Subsequently, beads were washed 3 × with PBS, and antibodies were eluted using 0.1 M Citric Acid (pH 2) and directly neutralized in 3 M Tris (pH 8). Purified elephant IgG antibodies were digested overnight using the Fab Preparation Kit (Pierce™) according to the manufacturer’s instructions. Following elution from the papain column, samples were incubated with pre-washed protein A/G Plus agarose beads (Pierce) for 6 hours on a roller mixer to remove undigested IgG. Beads were pelleted by centrifugation, and the supernatant containing unbound Fab fragments was collected.

Fab fragments were concentrated using Amicon Ultra filter columns and mixed with the purified gH/gL/gO complex at a 50–100-fold molar excess. The mixture was incubated overnight at 4 °C on a roller mixer to allow complex formation. The resulting gH/gL/gO–Fab complexes were separated from unbound Fab fragments and excess gH/gL/gO using an ÄKTA protein purification system (Cytiva). Fractions corresponding to the gH/gL/gO–polyclonal Fab peak were pooled and concentrated to ∼1 mg/mL prior to negative-stain electron microscopy analysis.

Subsequently, 3 µL of sample was applied to glow-discharged, carbon-coated copper grids (30 s, Cressington 208). After a 30 s adsorption period, grids were washed twice with Milli-Q water and stained with 2% (w/v) uranyl acetate. Grids were air-dried for ∼5 min and imaged on a Thermo Scientific Talos L120C transmission electron microscope operated at 120 keV and equipped with a CCD camera. A total of 208 micrographs were collected with a total dose of 30 e⁻/Å² at a magnification of 92,000×, yielding a final pixel size of 1.55 Å.

Data processing followed standard procedures similar to those outlined previously (51). Micrographs were imported into RELION (version 5.0.0) for processing. CTF parameters were estimated using CTFFIND (version 4.1), after which poor-quality micrographs were removed, resulting in a final dataset of 165 images. A Laplacian-of-Gaussian (LoG) picker was used for particle selection, producing an initial particle stack of 68,933 particles. Particles were extracted in a 240-pixel box and downsampled to 60 pixels, corresponding to a final pixel size of 6.2 Å. A single round of 2D classification was performed while ignoring CTF information until the first zero. Classes corresponding to gH/gL/gO or putative gH/gL/gO–polyclonal Fab complexes were retained, whereas classes representing overlapping particles or crowded fields were removed. This yielded a curated stack of 35,171 particles.

A 60 Å low-pass filtered version of the gH/gL/gO cryo-EM map was used as an initial model for 3D classification into 30 classes, again ignoring CTFs until the first zero. Particles belonging to a single well-defined 3D class consistent with a gH/gL/gO–Fab complex were selected and subjected to 3D auto-refinement.

### gH/gL/gO-cell binding assay

Primary Asian elephant umbilical cord endothelial cells (EUVEC; kindly provided by Petra van den Doel, ErasmusMC) were cultured in human endothelial serum-free medium (Gibco), supplemented with 20% heat-inactivated Fetal Bovine Serum (FBS), 100 ng/ml recombinant human Epidermal Growth Factor (EGF, Repligen) and 20 ng/ml recombinant human Fibroblast Growth Factor (FGF, Peprotech LTD). Culture flasks were coated with 20 µg/ml Bovine Fibronectin (Sigma Aldrich) prior to seeding of the cells. HEK293T cells were cultured in Dulbecco’s Modified Eagle’s Medium (DMEM, Gibco), supplemented with 10% FBS, 100 U/ml Penicillin and 100 µg/ml Streptomycin. Asian elephant PBMCs were isolated from heparinized whole blood samples using Histopaque®-1077 (Sigma-Aldrich®) within eight hours after blood collection and subsequently stored at -135°C. Samples were taken aseptically from ear or leg veins by zoo veterinary staff for diagnostic investigations according to standard veterinary practices after which their remainders were made available for the present study. Cells were gently thawed 1 day prior to analysis and incubated overnight in RPMI-1640 supplemented with 10% heat-inactivated FBS. All cells were maintained at 37°C and 5% CO_2_. Prior to use, adherent cells were harvested using enzyme-free cell dissociation buffer (Gibco), resuspended in buffer CI (1× PBS + 1 mM EDTA and 0.5% BSA) and kept at 4°C until further use. Per condition, 500.000 cells were used.

Excess biotin was removed from the purified gH/gL/gO and gH/gL protein samples using Zeba Spin desalting columns (Thermo Scientific) after which 2 µL streptactin-APC (IBA) was coupled to 1 μg protein in buffer CI (total reaction volume: 20 µL), and incubated in the dark for 30 min at 4°C. When indicated, the gH/gL/gO-streptactin-APC complex was incubated with serum or purified antibodies from naturally infected adult elephants for 1 hour at 4°C. The (preopsonized) APC-coupled protein was subsequently added to the dissociated cells and incubated in the dark for 30 min at 4°C. Cells were washed twice using 100 µl buffer CI, resuspended in 200 µl buffer CI and analyzed using the CytoFLEX LX Flow Cytometer (Beckman Coulter). Per condition 20.000 events were measured and data was analyzed using FlowJo software (v.10.8.1).

## Data Availability

Atomic coordinates will be deposited in the Protein Data Bank. The negative-stain and cryo-EM density maps will be deposited in the Electron Microscopy Data Bank, and the unaligned, non–gain-normalized movies will be deposited in the Electron Microscopy Public Image Archive.

Correspondence regarding structural data should be addressed to D.L.H., and requests regarding wet-lab data should be directed to T.E.H.

## Ethics statement

Ethical approval was not sought for the present study since it only made use of archival elephant samples taken under routine veterinary care by zoo veterinary staff and subsequently shared with our university for research purposes.

## Author Contributions

O.D.A.: Conceptualization, Investigation, Formal Analysis, Data Curation, Visualization, Writing – Original Draft

L.E.: Investigation, Formal Analysis, Visualization J.K.: Investigation, Formal Analysis

I.D.: Investigation, Resources S.Y.L.: Methodology

V.P.M.G.R.: Resources, Funding Acquisition W.S.: Resources, Funding Acquisition F.J.M.K.: Resources, Funding Acquisition

B.J.B. : Resources, Supervision, Funding Acquisition

C.A.M.H.: Conceptualization, Supervision, Project Administration, Funding Acquisition

D.L.H.: Conceptualization, Recourses, Formal Analysis, Data Curation, Visualization, Supervision, Project Administration, Funding Acquisition, Writing – Original Draft

T.E.H.: Conceptualization, Investigation, Formal Analysis, Data Curation, Visualization, Supervision, Project Administration, Funding Acquisition, Writing – Original Draft

All authors: Writing - Review & Editing

## Acknowledgements

We thank members of the Utrecht Virology Lab for valuable feedback during the preparation of this manuscript, and the staff of the Utrecht University Electron Microscopy Centre for their technical assistance. We would like to thank all zoos that provided elephant samples for this study for the collection and sharing of these samples.

