## Supplementary Figures and Tables for "The gH/gL/gO Complex of Elephant Endotheliotropic Herpesvirus 1A Functions as a Receptor-Binding Complex"

| Protein | Molecular weight[a] (kDa) | Predicted N-linked glycosalytion sites[b] | Expected molecular weight including glycans[c] (kDa) |
|---|---|---|---|
| gH-3×ST | 84.6 | 8 | 104.6 |
| gL-3×ST | 34.8 | 4 | 44.8 |
| gL-6×His | 29.9 | | 39.9 |
| gO-3×ST | 30.5 | 7 | 48.0 |
| gO | 24.2 | | 41.7 |
| PNGase F | 36.0 | - | - |

**Supplementary Table 1. Molecular weights of the protein constructs used in the study.** [a] Molecular weights (MWs) were predicted using the expasy PI/Mw tool (1). [b] The number of N-linked glycosylation sites were predicted using the NetNGlyc server (2). [c] The MW of glycosylated proteins was estimated by adding 2.5 kDa per predicted N-linked glycan[b] to the protein MW calculated in [a].

| Animal # | Age (y) | ΔOD | |
|---|---|---|---|
| | | gH/gL/gO − gH/gL | gO |
| 2 | 1,3 | 2,595 | 0,025 |
| 3a | 1,7 | 2,367 | 0,080 |
| 3b | 1,7 | 1,069 | 0,153 |
| 4a | 1,8 | 2,158 | 0,324 |
| 4b | 2,2 | 3,017 | 0,155 |
| 6a | 2,2 | 0,657 | 0,129 |
| 6b | 2,2 | 1,330 | 0,077 |
| 8 | 2,2 | 0,285 | 0,087 |
| 9 | 3,1 | 1,140 | 0,147 |
| 10b | 5,2 | 0,575 | -0,069 |
| 12 | 4,9 | 0,294 | 0,836 |
| 13b | 8,6 | 0,438 | 0,007 |
| 16 | 6,2 | 0,012 | -0,052 |

**Supplementary Table 2. Glycoprotein O-specific reactivity in sera of naturally infected Asian elephants.** Reactivity to gO out of the context of gH/gL/gO heterotrimer was determined using a gO-specific ELISA, whereas serum reactivity to gO in the context of the gH/gL/gO heterotrimer was determined by subtracting the ΔOD value obtained in the gH/gL ELISA from the ΔOD value obtained in the gH/gL/gO ELISA (gH/gL/gO − gH/gL). Only sera with non-saturated gH/gL-specific responses are shown. Longitudinal samples of individual elephants are identified by the individual elephant number followed by a character (a or b).

| | EEHV 1A gH/gL/gO (EMDB-XXXX) (PDB XXXX) |
|---|---|
| **Data collection and processing** | |
| Magnification | 165,000 |
| Voltage (kV) | 300 |
| Electron exposure (e–/Å$^2$) | 50 |
| Defocus range (μm) | -0.75 to -1.5 |
| Pixel size (Å) | 0.73 |
| Symmetry imposed | C1 |
| Initial particle images (no.) | 891,369 |
| Final particle images (no.) | 130,542 |
| Map resolution (Å) | 3.2 |
| FSC threshold | 0.143 |
| Map resolution range (Å) | 2.6-34.9 |
| **Refinement** | |
| Initial model used (PDB code) | NA |
| Model resolution (Å) | NA |
| FSC threshold | 0.143 |
| Map sharpening *B* factor (Å$^2$) | 110.9 |
| Model composition | |
| Non-hydrogen atoms | 17996 |
| Protein residues | 1097 |
| Ligands | 11 |
| *B* factors (Å$^2$) | |
| Protein | 41.89 |
| Ligand | 53.88 |
| R.m.s. deviations | |
| Bond lengths (Å) | 0.003 (0) |
| Bond angles (°) | 0.650 (0) |
| Validation | |
| MolProbity score | 1.85 |
| Clashscore | 8.82 |
| Poor rotamers (%) | 0.00 |
| Ramachandran plot | |
| Favored (%) | 94.41 |
| Allowed (%) | 5.59 |
| Disallowed (%) | 0.00 |

**Supplementary Table 3. Cryo-EM data collection, refinement and validation statistics.**

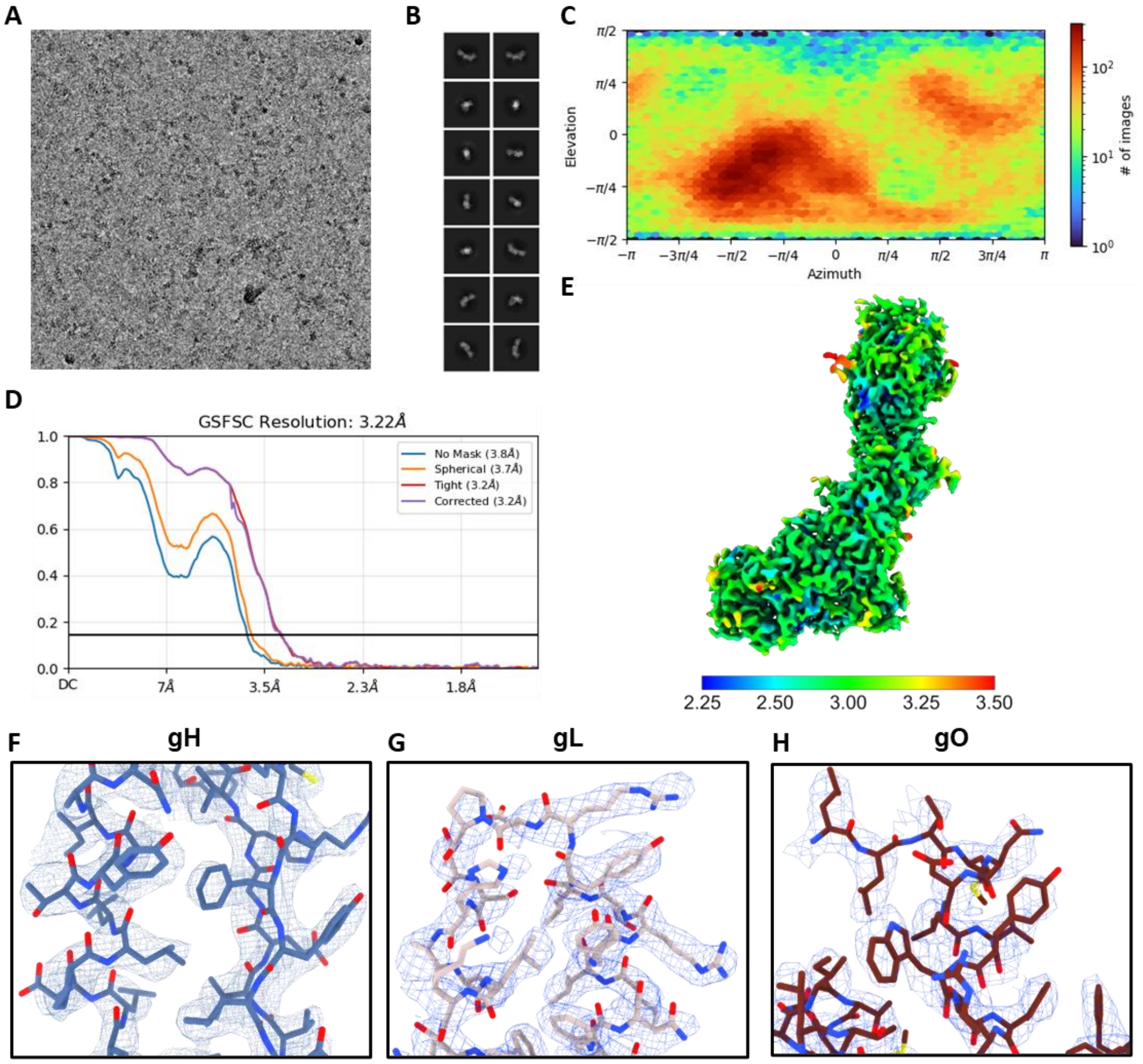

**Supplementary figure 1. Single-particle cryo-EM data processing for the gH/gL/gO complex**. (**A**) Representative micrograph. **(B)** Representative 2D classes for the gH/gL/gO complex. **(C)** Angular distribution plot of the final gH/gL/gO - C1 refined EM density map. **(D)** Gold-standard Fourier shell correlation (FSC) curve generated from the independent half maps contributing to the 3.2 Å global resolution density map of the the gH/gL/gO complex. (**E**) Local resolution filtered EM density map for the refined gH/gL/gO complex, colored according to local resolution which was calculated in CryoSPARC. (**F**) Zoomed-in view of gH occupying the according EM density, shown as a mesh. **(G)** Zoomed-in view of gL occupying the according EM density, shown as a mesh **(H)** Zoomed-in view of gO occupying the according EM density, shown as a mesh.

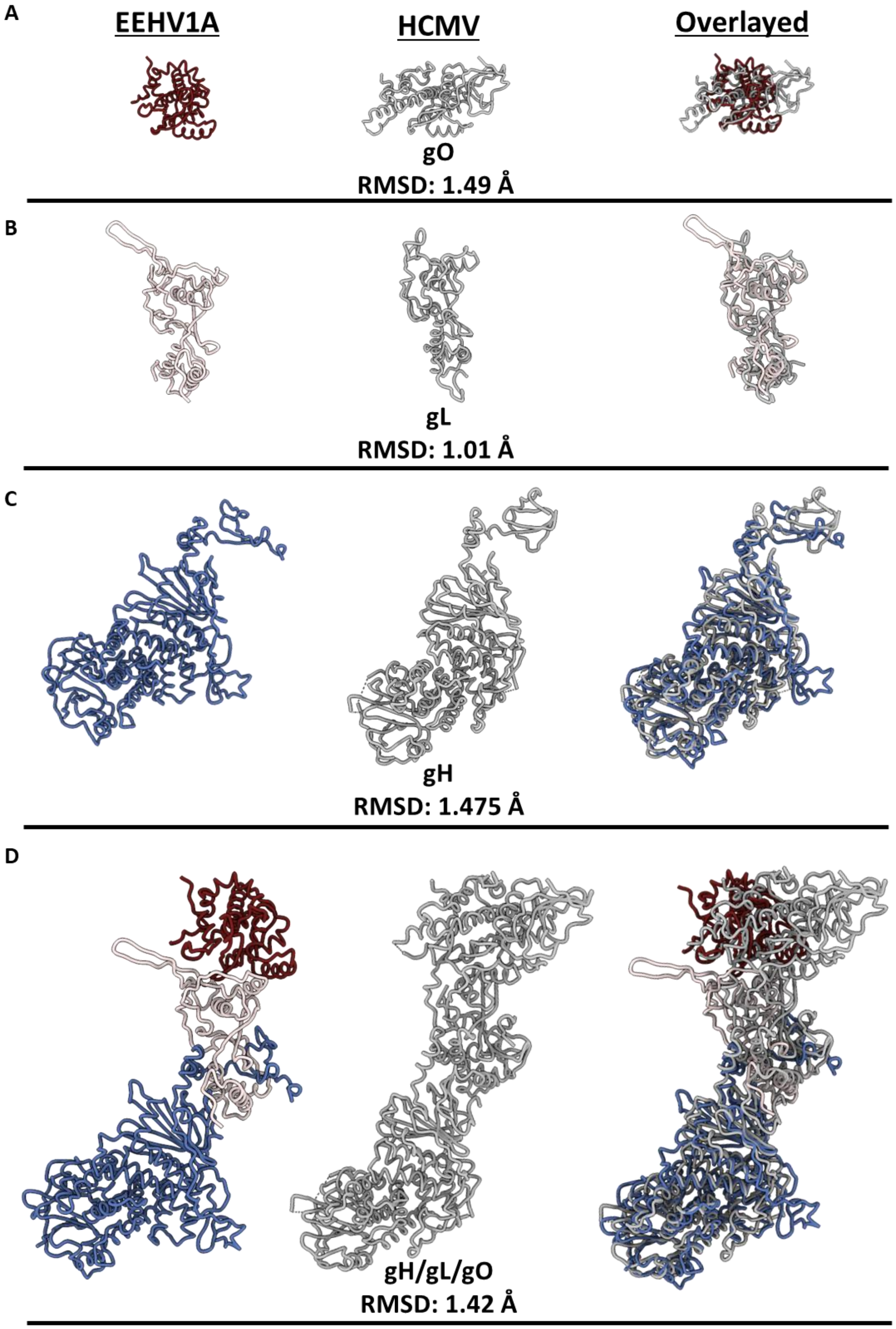

**Supplementary Figure 2. Structural comparison of EEHV1A gH, gL, and gO with homologous proteins from HCMV. (A)** Ribbon diagrams of EEHV1A gO (left), HCMV gO (middle), and their structural overlay (right). **(B)** As in panel A, shown for gL. **(C)** As in panel A, shown for gH. **(D)** Comparison of the complete

EEHV1A gH/gL/gO assembly (left) with the HCMV gH/gL/gO trimer (middle), and a superposition of both complexes (right). EEHV1A gH/gL/gO adopts a more compact, upward-oriented architecture compared with the more elongated HCMV trimer, while maintaining the conserved three-chain topology characteristic of betaherpesvirus gH/gL/gO complexes.

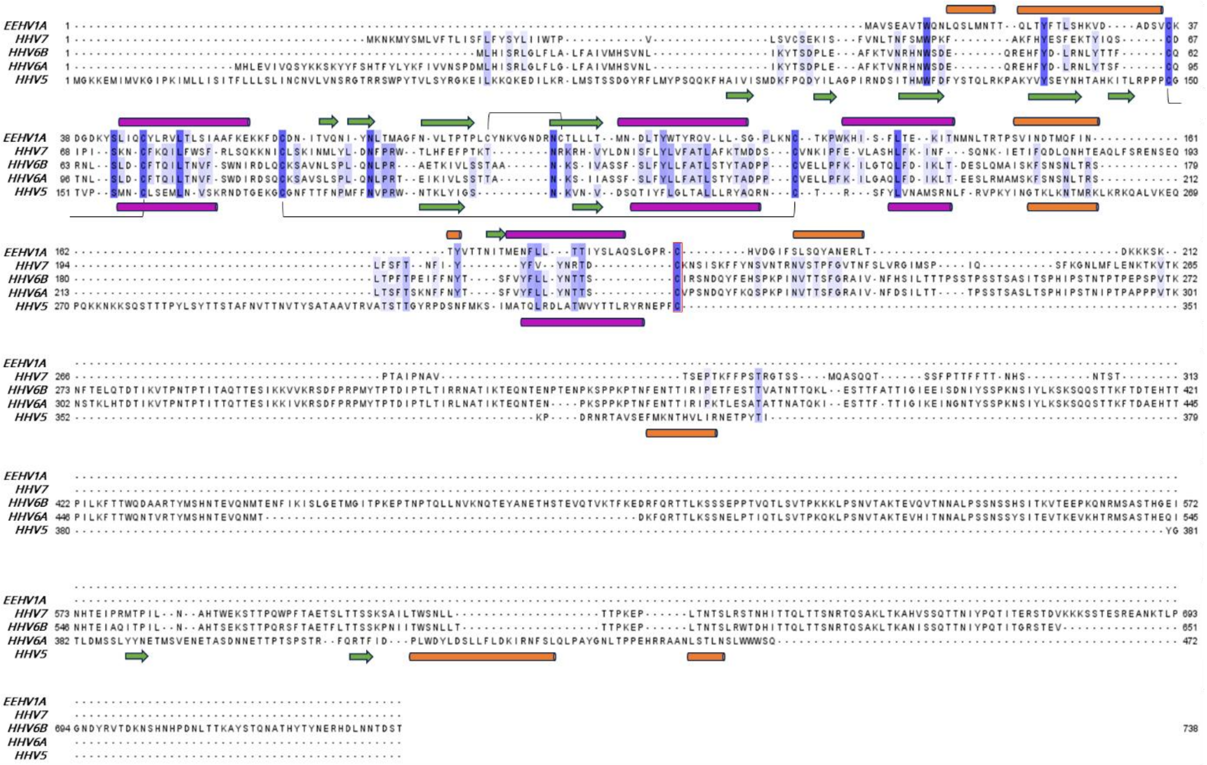

**Supplementary Figure 3. Amino acid alignment of EEHV1A and human betaherpesvirus gO.** Sequences (EEHV1A strain Kimba: AAG16085.1, HHV5 (HCMV) strain Merlin: YP_081522.1, HHV6A strain U1102: NP_042940.3, HHV6B strain Z29: NP_050228.1, HHV7 strain RK: AAC40761.1) were retrieved from GenBank, aligned in Jalview using the Clustal Omega webserver and subsequently edited manually to better align structural features that are conserved between HHV5 and EEHV gO, including cysteines, the alphahelices forming the cytokines fold and two beta sheets forming a scaffold associated with the cytokine fold. Sequences are coloured based on conservation (threshold 10%). For both EEHV1A and HHV5 (HCMV), alphahelices (cilinders), betasheets (arrows), and cysteine bonds (black lines) are indicated above (EEHV1A) respectively below (HHV5) the alignment. The alphahelices forming the cytokine fold are shown in purple, while alphahelices that are not part of the cytokine fold are shown in orange. The cysteines in the red frame (a.o. EEHV1A C189 and HCMV C351) are conserved between EEHV1A and human betaherpesviruses and form a disulfide bond with gL (EEHV1A C166 and HCMV C144).

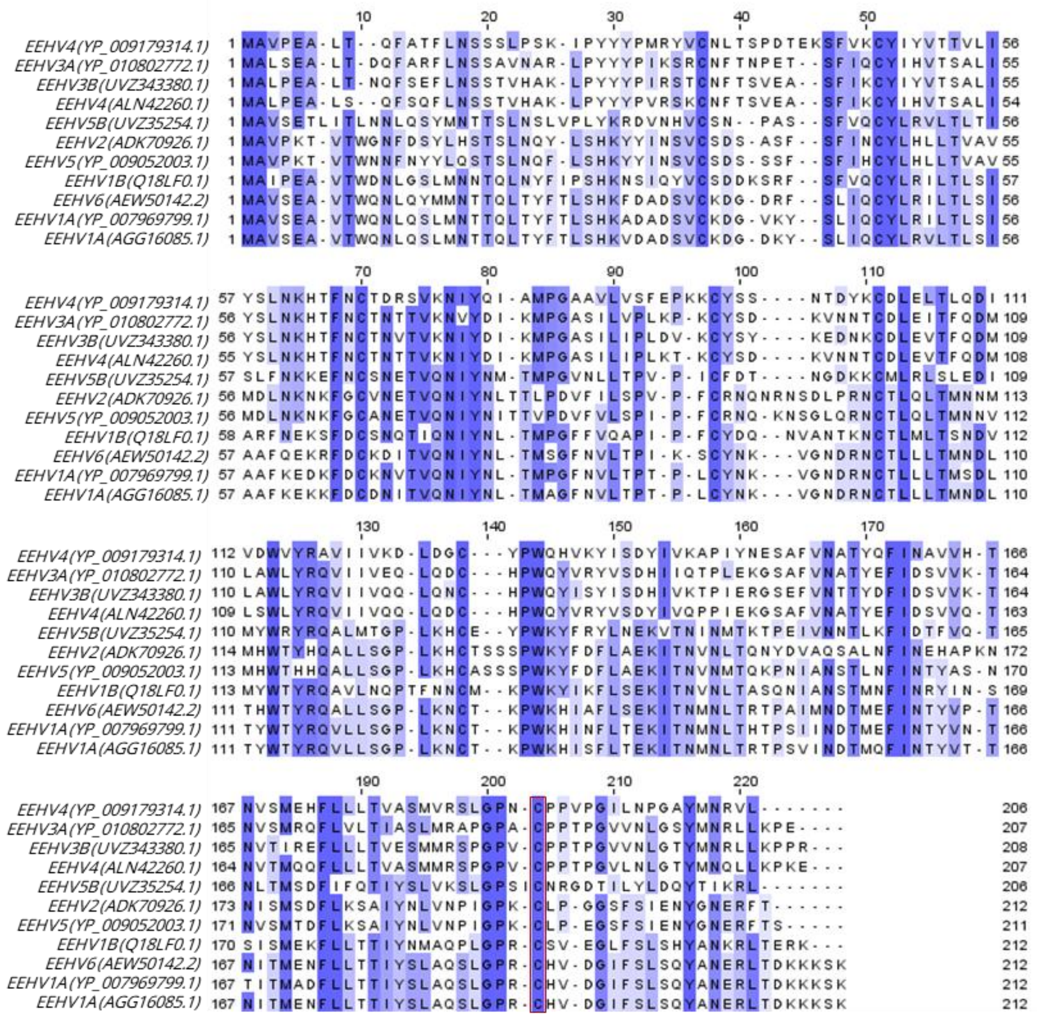

**Supplementary Figure 4. Amino acid alignment of glycoprotein O of various EEHV (sub)species.** Sequences (accession numbers indicated in the alignment labels) were retrieved from GenBank, aligned in Jalview using the MUSCLE webserver, and subsequently coloured based on conservation (threshold 10%). The cysteines framed in red are conserved across all EEHV (sub)species and correspond to EEHV1A gO-C189, the cysteine that forms a disulfide-linked bond with EEHV1A gL-C166.

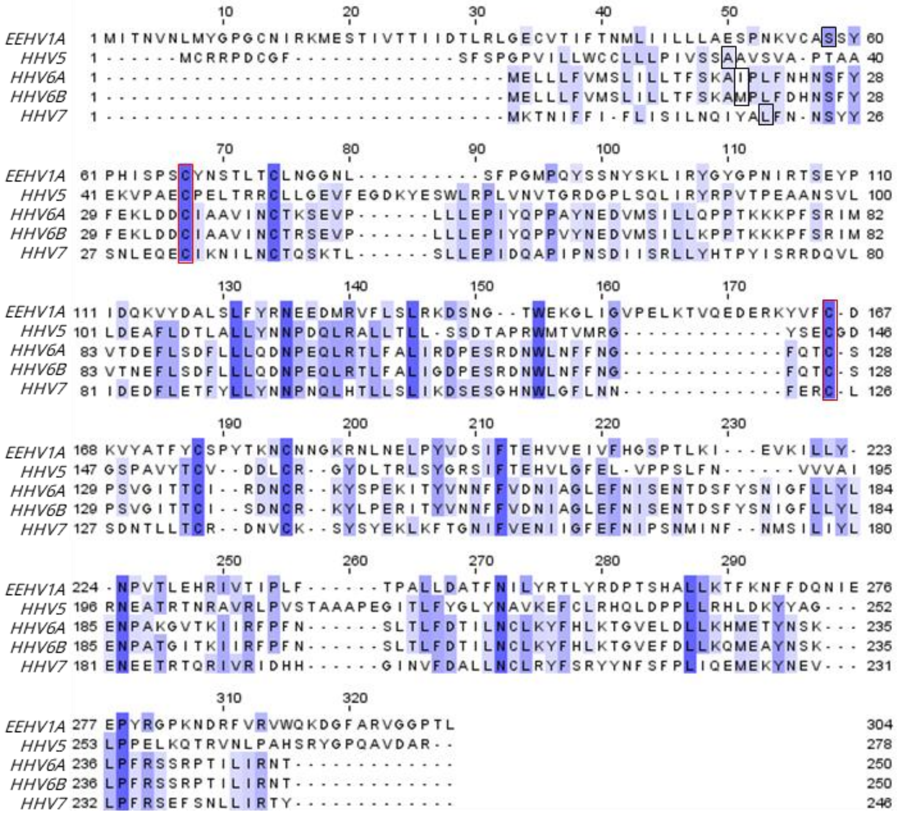

**Supplementary figure 5. Amino acid alignment of EEHV1A and human betaherpesvirus gL.** Sequences (EEHV1A strain Kimba: AGG16117, HHV5 (HCMV) strain Merlin: YP_081555, HHV6A strain U1102: NP_042975, HHV6B strain Z29: NP_050261, HHV7 strain RK: YP_073820) were retrieved from GenBank, aligned in Jalview using the MUSCLE webserver, and subsequently coloured based on conservation (threshold 10%). The amino acid residues shown in a black box indicate the first amino acid residue after the predicted signal peptide cleavage site (3). The cysteines indicated in the red box are conserved across human betaherpesviruses and EEHV1A and form disulfide-linked bonds with either gH (in conjunction with HCMV gL-C47 or EEHV1A gL-C67) or gO (in conjunction with HCMV gL-C144 or EEHV1A gL-C166).

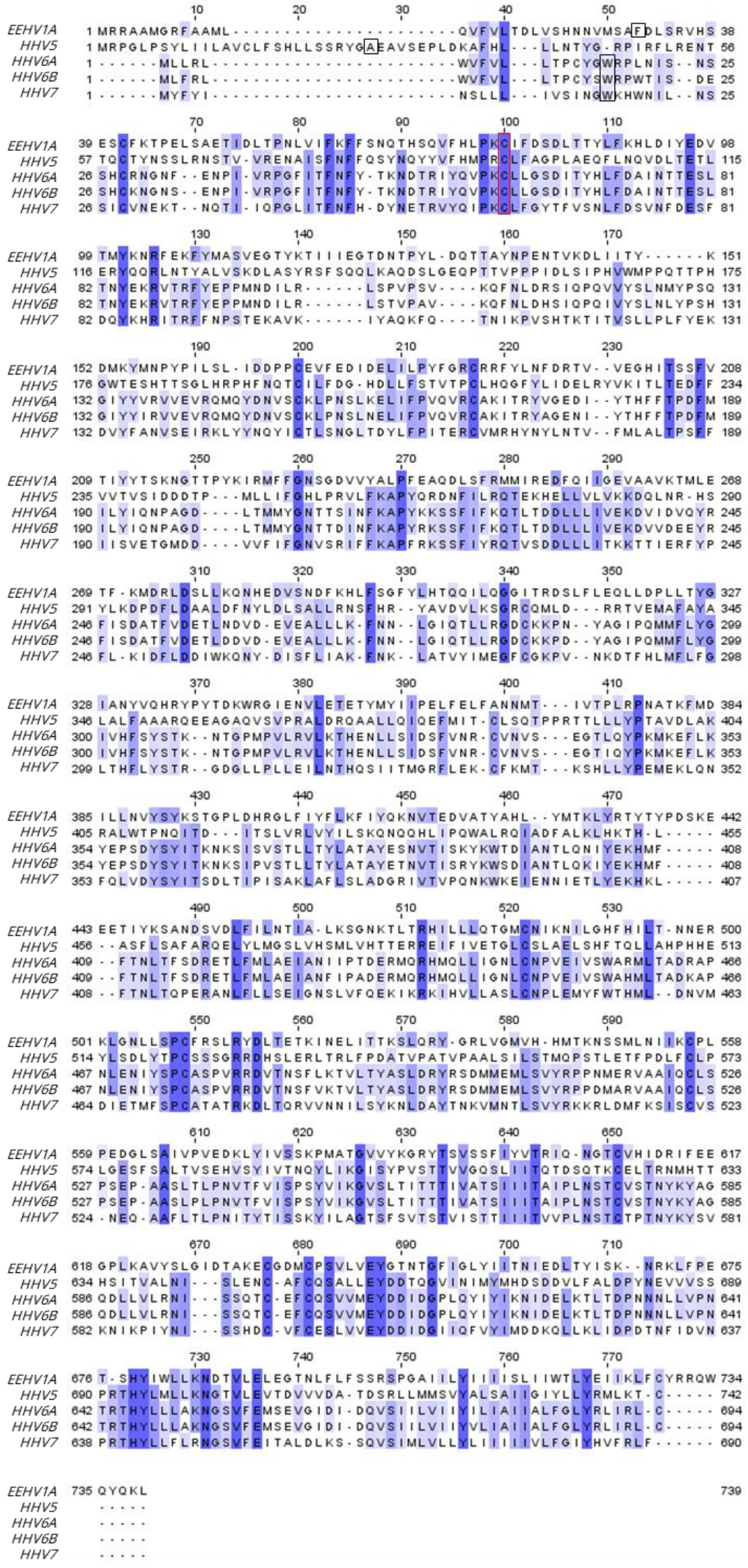
EEHV1A
HHV5
HHV6A
HHV6B
HHV7

**Supplementary figure 6. Amino acid alignment of EEHV1A and human betaherpesvirus gH.** Sequences (EEHV1A strain Kimba: AGG16086, HHV5 (HCMV) strain Merlin: YP_081523, HHV6A strain U1102: NP_042941, HHV6B strain Z29: NP_050229, HHV7 strain RK: YP_073788) were retrieved from GenBank, aligned in Jalview using the MUSCLE webserver, and subsequently coloured based on conservation (threshold 10%). The amino acid residues shown in a black box indicate the first amino acid residue after the predicted signal protein cleavage site (3). The cysteines indicated in the red box are conserved across betaherpesviruses and EEHV1A and form disulfide-linked bonds with gL (in conjunction with HCMV gH-C95 and EEHV1A gH-C78).

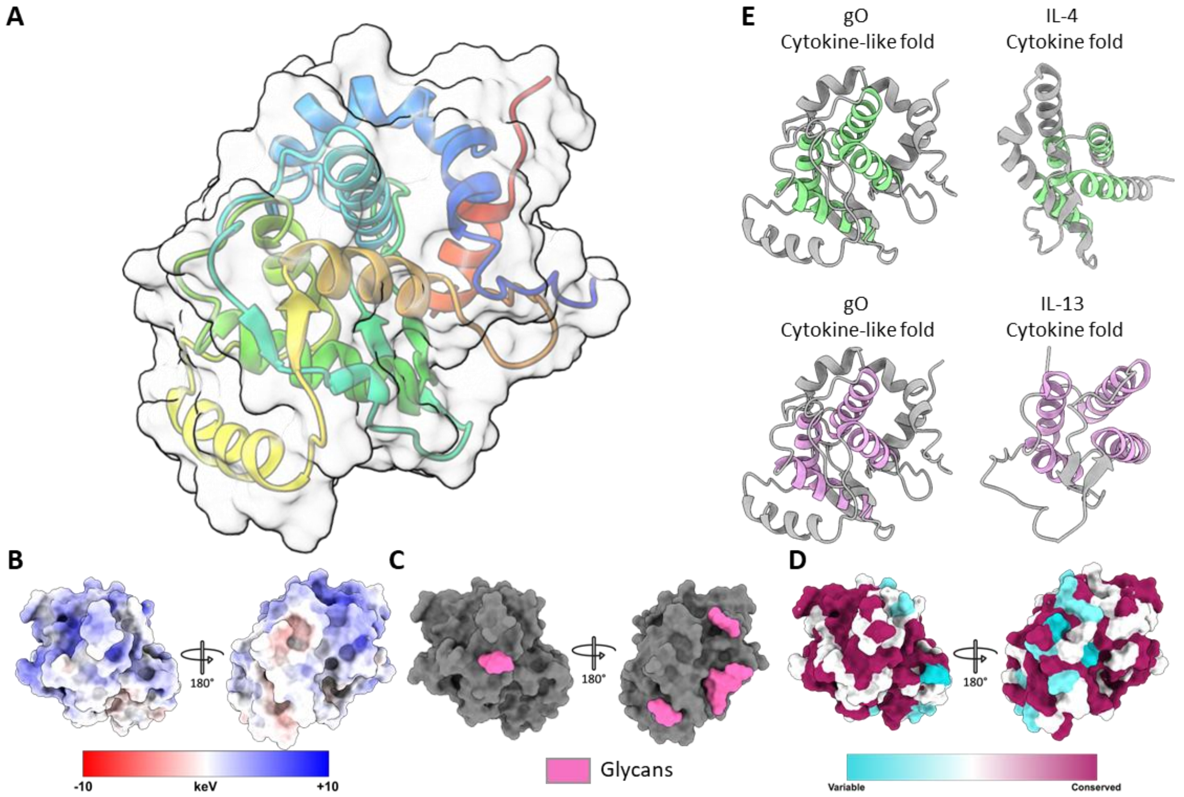

**Supplementary figure 7. The structure of the EEHV1A gO subunit (A)** Structure of the EEHV1A gO subunit with domain organization displayed and colored according to the chainbow coloring scheme. **(B)** Electrostatic surface of EEHV1A gO subunit. **(C)** Glycosylation site distribution of the EEHV1A gO subunit. **(D)** Conservation surface of EEHV1A gO subunit based on sequences from EEHV1A strains. **(E)** Secondary structure comparison between core folds of EEHV1A gO subunit and cytokine folds of both IL-4 and IL-13 (PDB: 2B8U and PDB: 1IJZ). IL-4 like folds highlighted in Chartreuse and IL-13 folds highlighted in pink.

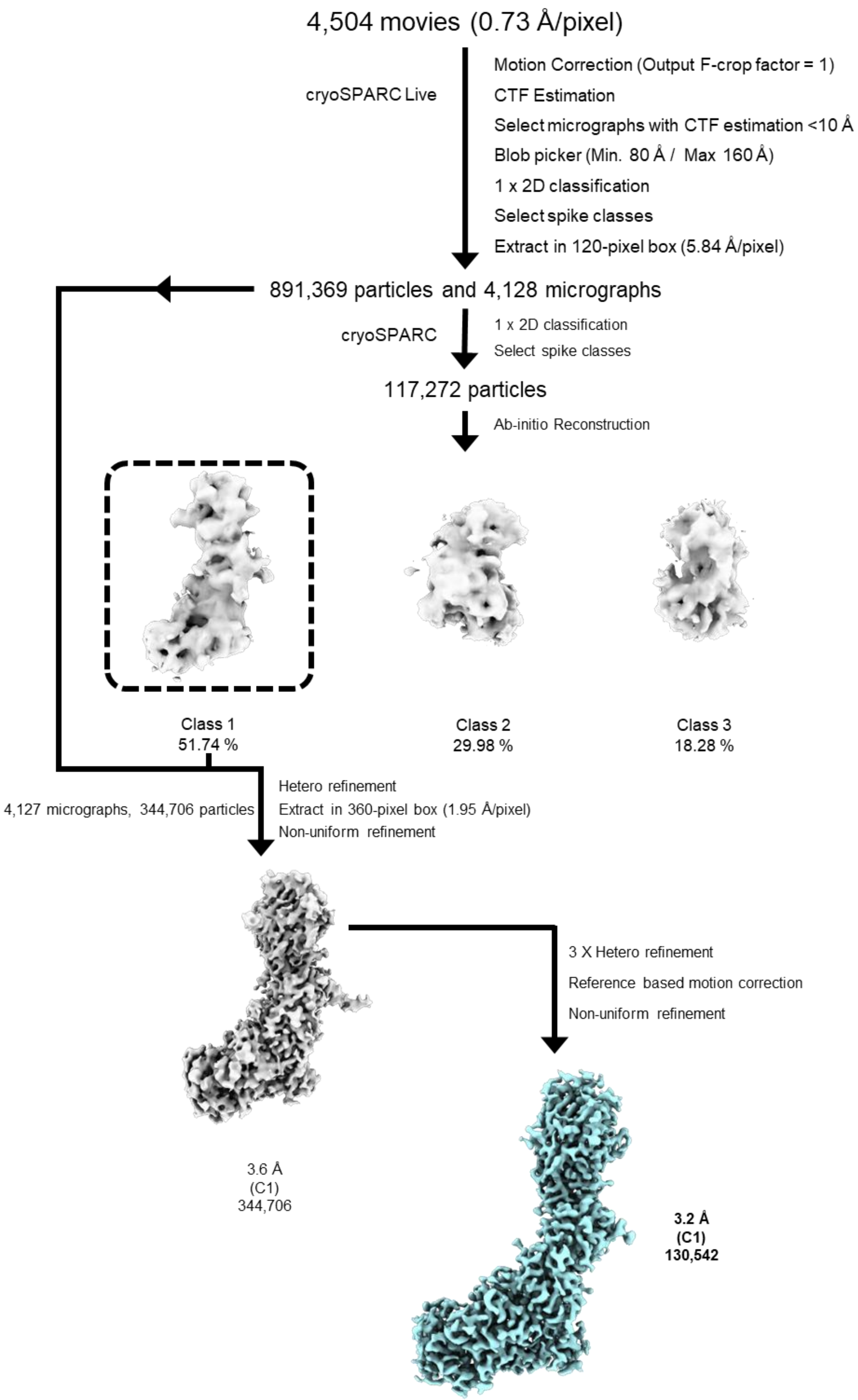

**Supplementary Figure 8. Single-particle cryo-EM processing pipeline for the EEHV1A gH/gL/gO complex.** Overview of the cryo-EM workflow used to obtain the 3.2 Å reconstruction of the gH/gL/gO heterotrimer. Motion-corrected and CTF-estimated micrographs were used for automated particle picking, followed by iterative 2D classification and ab initio reconstruction. Subsequent rounds of heterogeneous refinement, particle re-extraction, and non-uniform refinement in cryoSPARC yielded the final high-resolution map. Particle numbers and intermediate reconstructions are shown at each step.
